# Gasdermin C links nutrient and immune signaling to protist-induced type 2 immunity and intestinal repair

**DOI:** 10.1101/2025.09.24.678435

**Authors:** Andrea Keller, Ayobami Adeniyi, Kübra B. Akkaya-Colak, Maria Festing, Andrew Schwieters, Rathan Kumar, James Sledziona, Khushboo Gupta, Patrick Stevens, Maciej Pietrzak, Vincenzo Coppola, Parvathi Ranganathan, Brian Ahmer, Adriana Forero, Sue E. Knoblaugh, Maria M. Mihaylova

## Abstract

The Gasdermin family of proteins has recently been implicated in tissue repair and homeostasis through their effector function in type 2 immunity and pyroptosis. Yet the role of Gasdermin C proteins has not been fully characterized in the mammalian intestine, where environmental factors can influence epithelial regeneration and repair. Here we report that *Gsdmc2-4* genes are regulated in a nutrient-dependent manner and are suppressed with aging. We uncover that commensal protists in the gut regulate *Gsdmc2-4* expression through activation of type 2 immune responses. In intestinal organoid experiments, we find that STAT6 is necessary for *Gsdmc2-4* induction in response to type 2 cytokines; however, basal expression of *Gsdmc2-4 in vivo* is only partially diminished in *Stat6* knockout animals. Finally, in protist-colonized animals, loss of *Gsdmc1-4* exacerbated mucosal erosion and inflammation in response to Dextran sodium sulfate (DSS) exposure, implicating these proteins in coordinating epithelial responses to injury.

## INTRODUCTION

The intestine is one of the most regenerative organs in the body and one of the most dynamic environments. Perhaps due to constant exposure to a milieu of nutrients and byproducts of the microbiome, the entire intestinal epithelium is replaced every 4-5 days^1^. This process is supported by dedicated tissue-resident stem cells that differentiate into highly proliferative progenitor cells located in the upper isthmus of the intestinal crypts. Cells residing in this transit-amplifying region of the epithelium undergo rapid rounds of cell division prior to differentiating and committing to either a secretory lineage (e.g., goblet cells, tuft cells, enteroendocrine cells) or an absorptive lineage (enterocytes). After this point, cells continue to migrate up the villus length over the course of several days and are eventually shed off into the lumen – a process sometimes referred to as the “intestinal escalator”. Host epithelial cells also undergo complex interactions with commensal microbes, as well as resident lymphocytes, to maintain homeostasis. The intestine is responsible for more than just nutrient sensing and forms a barrier against pathogens through coordinated immune responses and repair of tissue damage. Specialized cell types contribute to each of these functions throughout the crypt and villus, and this balance can become greatly disrupted during aging, injury, or infection, resulting in changes to the composition and function of these cells, including decreased numbers of stem cells^2^, skewed differentiation to secretory cells^3,4^, and decreased gut barrier integrity^5^. Ultimately, an imbalance in the epithelial barrier function can lead to microbial invasion, inflammation, and progressive gut dysbiosis.

Gasdermin proteins are commonly expressed at epithelial surfaces and in immune cell types, and they are known for their ability to protect against intracellular and extracellular pathogens through induction of pyroptosis^6^. Most members of the Gasdermin family (GSDMA, B, C, D, and E) are thought to be cleaved by inflammatory caspases to release an N-terminal fragment that self-oligomerizes and forms a pore in the cell membrane. This traditionally serves two purposes: to kill the infected cell, and to release cytokines and amplify pro-inflammatory signals. Extensive work has been done on GSDMD, primarily in immune cells, to elucidate the various pathogen and danger signals that can activate pyroptosis, to determine which caspases can cleave GSDMD, and to characterize the precise structures of the full-length protein, the cleaved protein, and the pore itself ^7,8,9,10,11,12^. Similarly, GSDMB and GSDME have been shown to be cleaved by caspases-3 and 7 to induce pyroptosis^13,14,15^. It is currently unknown whether GSDMA is cleaved by any caspases, but the N-terminal domain is capable of binding membrane phospholipids, as well as cardiolipin found in mitochondrial and bacterial membranes and can induce pyroptosis when overexpressed^16,7^. Until recently, very little was known about Gasdermin C (GSDMC), which is almost exclusively expressed in the gastrointestinal tract and skin. Humans have one copy of the gene (*GSDMC*), while mice have four (*Gsdmc1*, *Gsdmc2*, *Gsdmc3*, *Gsdmc4*), of which the latter three are most highly expressed.

Recently, the GSDMC family of proteins has gained attention for its potential role in type 2 immune responses against helminths in the intestine^17,18^, as well as possible anti-tumor pyroptotic effects in cancer cells^19,20^. New studies show that murine *Gsdmc2-4* are upregulated following infections with either of the parasitic worms *Nippostrongylus brasiliensis* or *Heligmosomoides polygyrus*. These large extracellular pathogens are sensed by specialized epithelial cells in the intestine called tuft cells, which secrete IL-25 to activate type 2 innate lymphoid cells (ILC2s). In turn, ILC2s secrete IL-4 and IL-13 to promote differentiation of intestinal stem and progenitor cells towards secretory cell types like goblet cells that will produce mucus to capture the parasitic worm and eliminate it, a process often called the “weep and sweep” response^21–24^. However, what is less apparent is if pyroptosis plays a role in the induction of this type 2 immune response and if GSDMC is indeed creating pores for the purpose of inducing massive cell death in the intestine. It is also not well understood how Gasdermin C proteins respond to other commensal organisms under homeostatic conditions, in particular protists, which have been shown to elicit a type 2 immune response^22,25^.

Finally, how environmental factors such as diet or age impact host defense through GDMCs needs to be further evaluated. Here, we discovered that *Gsdmc* transcripts were regulated in response to nutrient cues and were suppressed with age. We also found, like recent reports, that GSDMC proteins were strongly induced by type 2 immune cytokines IL-4 and IL-13 in intestinal epithelial cells. Importantly, we established that these signals are altered by the intestinal microenvironment and that *Gsdmc1-4* expression under homeostatic conditions is dependent on commensal intestinal microbial presence, which varies between facilities. In addition, loss of *Gsdmc1-4* in protist-colonized animals increases DSS induced inflammation and mucosal erosion, suggesting a potential role in epithelial repair. Collectively, these findings reveal that GSDMC proteins can act as nutrient- and age-sensitive regulators of intestinal epithelial responses, integrating environmental and immune signals to promote tissue protection and repair.

## RESULTS

### Gasdermin 2-4 are enriched from proximal to distal end of intestine and responsive to nutrient abundance

Using our previous bulk RNA sequencing analysis^2,26^ on isolated small intestinal stem and progenitor cells from either mice under *ad libitum* (AL) or fasting conditions, we discovered that *Gsdmc2-4* were highly enriched in intestinal progenitor cells^2^. To corroborate these findings, we isolated small intestinal (SI) crypts, enriched for progenitor cells and found by RT-qPCR that *Gsdmc2*, *Gsdmc3*, and *Gsdmc4* are abundantly expressed, whereas *Gsdmc1* is largely absent (Fig. 1A). Accordingly, our studies focused on *Gsdmc2-4*. We further noted that *Gsdmc2-4* transcripts and protein levels increased from the proximal to distal end of the SI, with their expression being the highest in the colon (Fig. 1A, 1B, S1A, S1B). In addition, *Gsdmc2-4* transcripts were primarily expressed in progenitor cells located at the top of the crypts/bottom of the villi in the mouse small intestine, and in a similar zone of transit-amplifying cells in the colon (Fig. 1B, upper panel), which has also recently been reported^18^. Moreover, we found that GSDMC2/3 proteins are expressed in both the upper crypt and in villi epithelial cells (Fig. 1B, lower panel, and S1B). The expected full-length GSDMC2/3 protein was detected in small intestinal and colon crypt lysates, as well as villi lysates, with the highest abundance found in distal SI villi and the colon (Fig. S1A, S1B). Smaller protein products were also detected and increased in abundance in proportion to the full-length protein (Fig. S1A, S1B). Collectively, these observations suggest that *Gsdmc* transcripts are largely produced in progenitor cells residing in the transient amplifying zone, while the protein is produced and remains stable in these cells as they differentiate and migrate up the villus over several days. In contrast, *Gsdmd* transcript and protein were present primarily in mature enterocytes and mature colonocytes (Fig. 1B).

**Figure 1:**
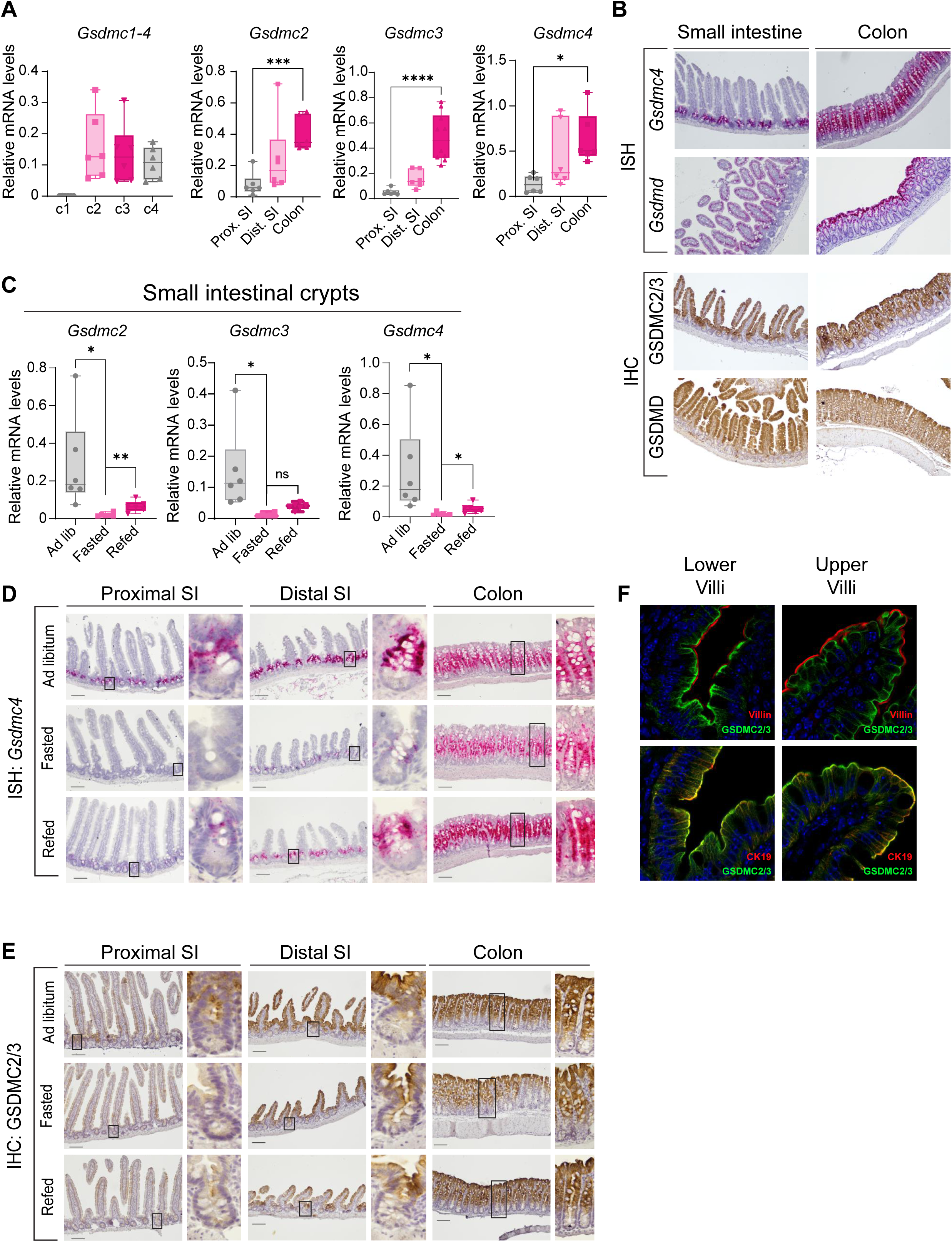
Gasdermin C2-4 levels change in response to nutrient availability. A) *Gsdmc1*, *Gsdmc2*, *Gsdmc3*, or *Gsdmc4* expression were evaluated by RT-qPCR analysis in either whole small intestine (SI) crypts or crypts collected from proximal SI, distal SI, and colon (n=6). (B) RNAscope *in situ* hybridization staining of either *Gsdmc4* or *Gsdmd* and IHC staining of either GSDMC2/3 or GSDMD in mouse small intestine and colon. (C) SI crypts from *ad libitum* (n=6), 24hr fasted (n=6), and 24hr refed (n=6) mice were collected and expression of *Gsdmc2*, *Gsdmc3*, and *Gsdmc4* genes were determined by RT-qPCR. (D) RNAscope *in situ* hybridization staining of *Gsdmc4* in proximal and distal regions of mouse small intestine and in whole colon from *ad libitum* (n=3), 24hr fasted (n=3), and 24hr refed (n=3) mice. (E) IHC staining of Gsdmc2/3 in proximal and distal regions of mouse small intestine and in whole colon from *ad libitum* (n=3), 24hr fasted (n=3), and 24hr refed (n=3) mice. (F) Immunofluorescent staining of mouse small intestine for GSDMC2/3 (green) and villin (red) or cytokeratin-19 (CK19 in red). Data are presented as mean ± SD. *p < 0.05, ***p<0.001, and ****p<0.0005.

Re-analyzing further our published RNAseq data^2,26^, we noted that *Gsdmc2-4* are among of the most downregulated genes in intestinal progenitor cells following a 24-hour fasting compared to *ad libitum* controls. We followed this up with RT-qPCR analysis of isolated small intestinal crypt cells (Fig. 1C), as well as *in situ* hybridization (Fig. 1D). Following 24-hour nutrient deprivation (fasting), *Gsdmc2-4* transcript levels rapidly declined, as seen by near-complete loss of staining in the transit-amplifying region of the small intestine and a decrease in signal intensity in the colon (Fig. 1D). Upon refeeding, transcript levels in previously fasted mice recovered (Fig. 1D). Assuming the primary factor affecting GSDMC2/3 protein expression is transcript availability and that any current protein will persist over the course of several days as epithelial cells migrate up the villus, any changes in protein levels may thus require longer to become detectable. Indeed, small intestinal villi showed a slight decrease in GSDMC2/3 protein between *ad libitum* and 24- hour fasted mice, an effect which was more apparent in fasted crypts (Fig. 1E), suggesting that the upper crypt cells are where the transcript is likely being regulated. Following 24-hour refeeding, we still detected an overall slight decrease in protein expression in the villi (Fig. S1C). Interestingly, we observed a region of low GSDMC2/3 protein levels mid-villus, where clusters of cells had begun to migrate up the villi that had experienced the 24-hour nutrient deprivation and consequently produced less of the *Gsdmc2/3* transcript and protein. Older cells that had already produced GSDMC2/3 protein prior to the fasting period had arrived at the villus tip and had higher expression of GSDMC2/3; newer cells that were produced following the refeeding had normal GSDMC2/3 expression closer to the progenitor zone as well (Fig. 1E). *Gsdmc2-4* expression levels did not change significantly in the colon, indicating the colon is less sensitive to changing nutrient availability (Fig. 1E, S1D, and S1E). In contrast, *Gsdmd* expression remained largely unaffected by nutrient status (Fig. S1F). This provides an interesting scenario where dietary fluctuations may selectively perturb *Gsdmc2-4* levels and thus potentially affect gut barrier homeostasis.

We next sought to define the subcellular localization of GSDMC2/3 as preliminary staining results indicated that it may be expressed at, or near, the plasma membrane of epithelial cells, particularly in mature enterocytes in the villi. Co-staining with the brush-border protein villin showed that GSDMC2/3 is expressed just below the microvilli in epithelial cells (Fig. 1F). Further investigation into the cytoskeletal network beneath the plasma membrane led us to examine keratin-19 (CK19), a component of the terminal web of filaments located beneath the brush border of epithelial cells^27^. GSDMC2/3 colocalized with CK19 at the terminal web in both upper crypts and villi (Fig. 1F) but not with Villin. Together, these findings demonstrate that GSDMC2-4 exhibit region-specific and nutrient-dependent expression patterns in the intestine, distinct from GSDMD, highlighting their unique spatial distribution and regulatory dynamics relative to other Gasdermin family members.

### *Gsdmc2-4* expression is reduced in the aging microenvironment of the intestine

We previously took an unbiased approach to identify pathways and processes deregulated in aged animals and discovered that *Gsdmc2-4* were highly downregulated with aging in progenitor cells (Fig. 2A)^2^. We confirmed the decrease in *Gsdmc2-4,* primarily in transit-amplifying progenitor cells, via *in situ* hybridization staining in small intestinal and colon crypts (Fig. 2B), and we noted a trend towards lower *Gsdmc2*-*4* transcript levels in small intestinal crypts in old mice compared to young, as shown by RT-qPCR analysis (Fig. 2C and S2A). Expression of *Gsdmc2* in colon crypts was significantly downregulated in old mice, while *Gsdmc3* and *Gsdmc4* did not exhibit a strong difference (Fig. S2B). In contrast, transcript levels of *Gsdmd* were unchanged in small intestinal crypts between young and old mice (Fig. S2C). Protein levels of GSDMC2/3 also declined in the intestines of old mice consistently across crypts and villi, indicating a generalized reduction of Gasdermin C in the gut with aging (Fig. 2D and 2E).

**Figure 2:**
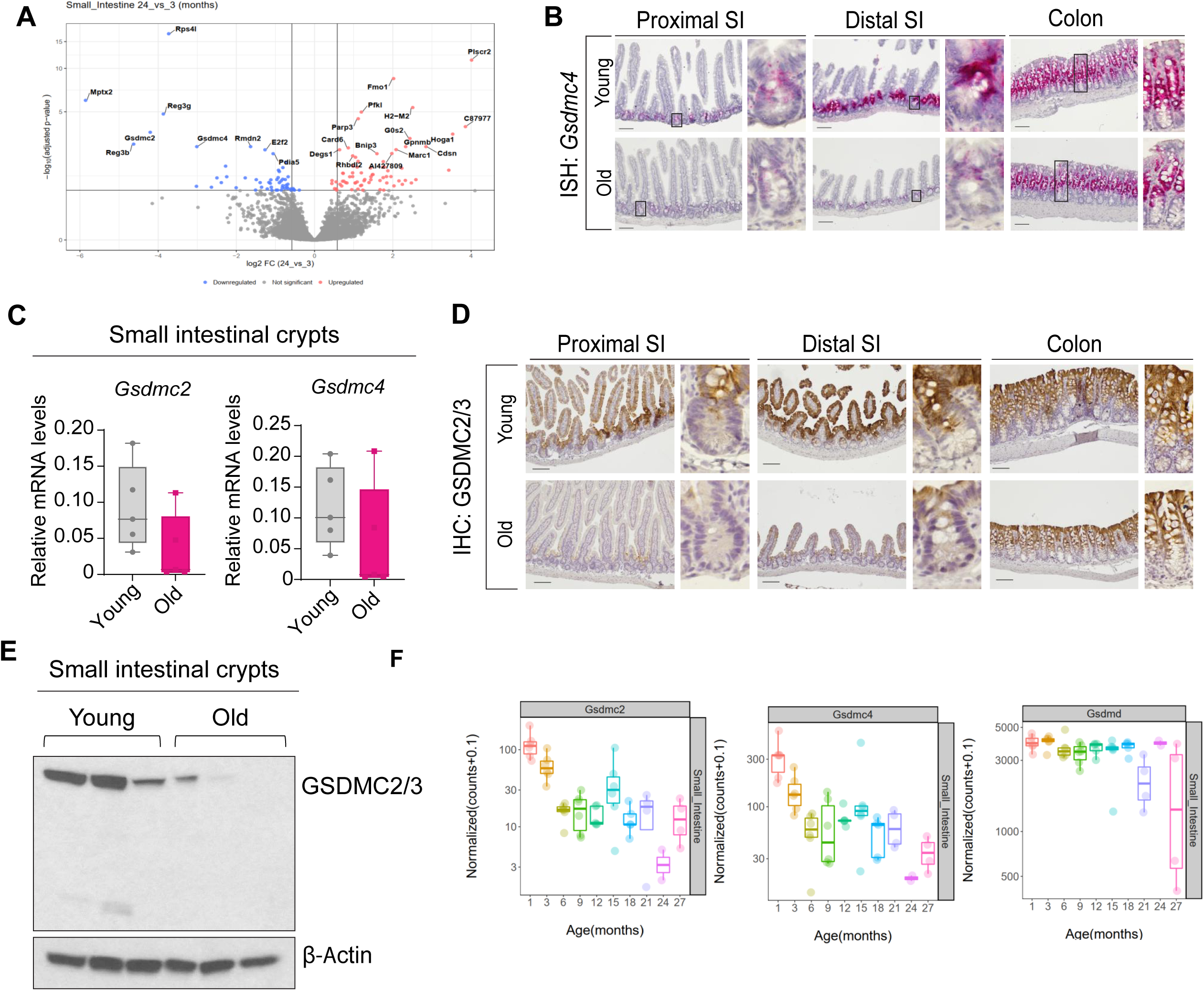
Expression of Gasdermin C2-4 decrease in aged mice. (A) Upregulated and downregulated genes in sorted intestinal progenitor (Lgr5-GFP^low^) cells from young and aged animals. (B) RNAscope *in situ* hybridization staining of *Gsdmc4* in proximal and distal regions of mouse small intestine and in whole colon from young (3-6 months, n=6) and aged (22-30mo, n=4) mice. (C) Whole SI crypts from young (3-6 months, n=5-6) and old (22-30mo, n=5-6) mice were collected and expression of *Gsdmc2*, *Gsdmc3*, and *Gsdmc4* genes were determined by RT-qPCR. (D) IHC staining of GSDMC2/3 in proximal and distal regions of mouse SI and in whole colon from young (3-6mo, n=6) and aged (22-30mo, n=4) mice. (E) Lysates from SI crypts from young (3-6mo, n=5-6) and old (22-30mo, n=5-6) mice were collected and immunoblotted for Gsdmc2/3. Representative samples are shown. (F) Box and whisker plots of normalized read counts from the Tabula Muris Consortium^27^ for *Gsdmc2*, *Gsdmc4*, and *Gsdmd* across the mouse lifespan.

To further validate our findings, we analyzed data available from the Tabula Muris Consortium^28^, a multi-organ compendium of murine transcriptional profiles throughout 27 months of the mouse lifespan. *Gsdmc2* and *Gsdmc4* were highly downregulated in the small intestine of 24-month-old versus 3-month-old mice, further confirming our findings (Fig. 2F). Additionally, there was a decline in *Gsdmc2-4* expression over the entire lifespan, while *Gsdmd* expression remained stable throughout life (Fig. 2F). Thus, GSDMC2-4 were selectively reduced in intestinal epithelium of aged mice while GSDMD remained unaffected, highlighting a potential Gasdermin C–specific vulnerability to age-associated shifts in tissue homeostasis.

### Type 2 immune signals regulate GSDMC protein levels in *ex vivo* organoid models

To investigate the possibility of immune signals regulating *Gasdermin C* expression, we purchased NOD-SCID mice^29^ and discovered that *Gsdmc2-4* transcript and protein expression were lower in the small intestinal crypts and villi of immunocompromised mice compared to wild-type mice (Fig. 3A and Fig. 3B). GSDMC2/3 protein levels were consistent between WT and NOD-SCID mice in the colon, while transcript levels were elevated in the immunocompromised mice (Fig. S3A). Using small intestinal crypts to generate organoids, we also found that *Gsdmc2-4* expression is practically undetectable in organoid cultures (Fig. 3C), further indicating that baseline expression of *Gsdmc2/4* in intestinal epithelial cells may be dependent on signals from the microenvironment, such as the microbiota or resident immune cells, two components that are absent from traditional organoid cultures.

**Figure 3:**
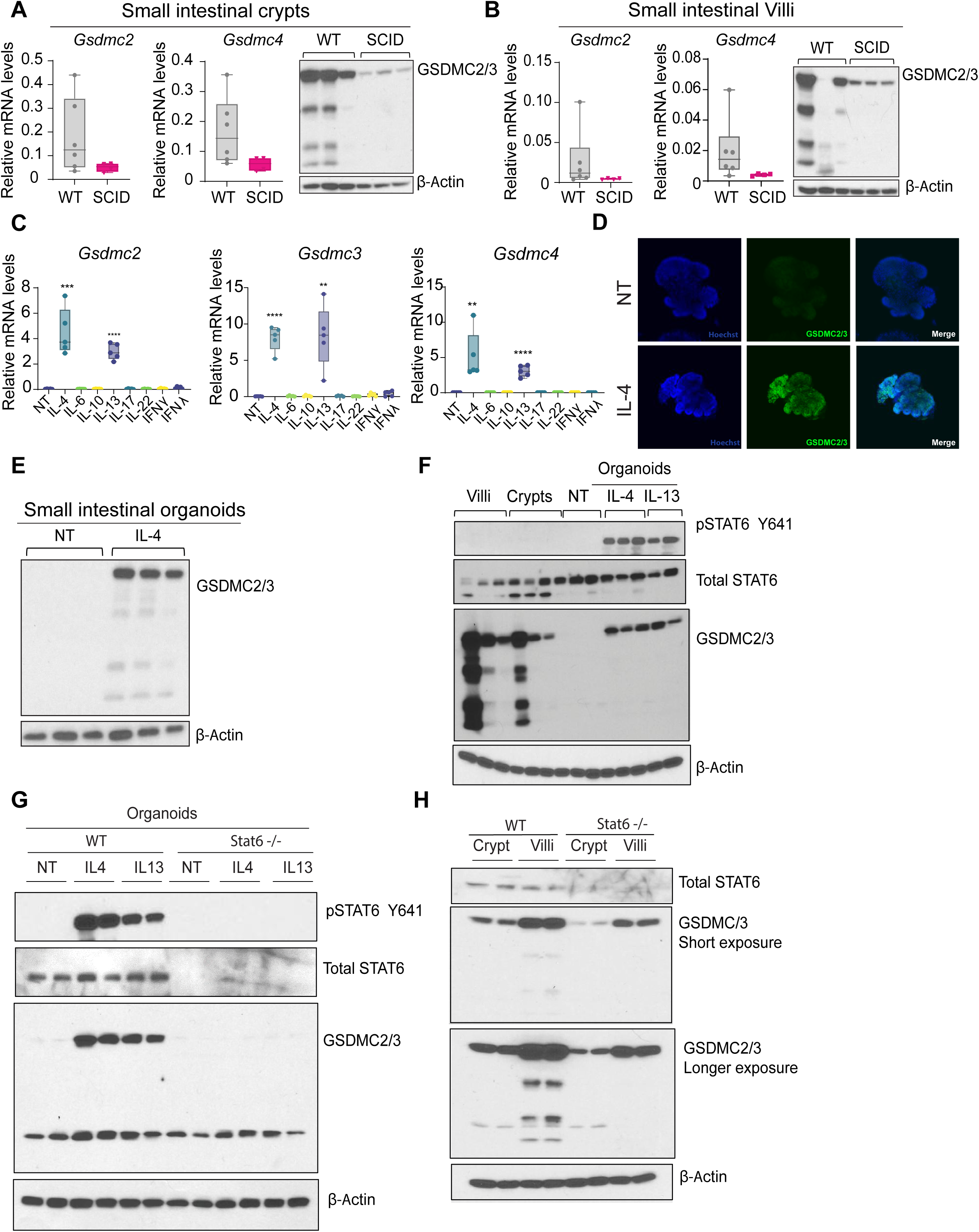
Type 2 immune signals upregulate Gasdermin C2-4. (A) SI crypts and (B) SI villi from WT (n=5-6) and NOD-SCID mice (n=4) were collected and used for RT-qPCR analysis of *Gsdmc2* and *Gsdmc4* and immunoblotted for GSDMC2/3. Representative samples are shown. (C) SI organoids were cultured for 4 days and treated with cytokines for 48hrs. Expression of *Gsdmc2*, *Gsdmc3*, and *Gsdmc4* genes was determined by RT- qPCR (n=5 biological replicates). (D) SI organoids were cultured for 4 days and treated with cytokines for 48hr, then fixed and stained for GSDMC2/3 (green). (E) In parallel, lysates were collected and immunoblotted for GSDMC2/3. Representative images and blots of n=5 biological replicates are shown. (F) SI organoids from WT (n=4) and *Stat6^-/-^* (n=4) mice were established and treated with IL4 and IL13 for 6hrs. Lysates were collected for immunoblotting. (G) Lysates from SI villi, SI crypts, and SI organoids treated with cytokines for 6 hours were collected for immunoblotting (n=3 biological replicates). (H) Lysates from SI crypts and villi from WT (n=4) and *Stat6^-/-^* (n=4) mice were collected for immunoblotting. Representative samples are shown. Data are presented as mean ± SD. **p < 0.01, ***p<0.001, and ****p<0.0005.

In order to more broadly examine what immune signals may regulate Gasdermin C expression, we treated small intestinal organoid cultures with a panel of cytokines, including IL-4, IL-6, IL-10, IL-13, IL-17, IL-22, IFNγ, and IFNλ and found that only the type 2 cytokines IL-4 and IL-13 were able to induce *Gsdmc2-4* expression (Fig. 3C and Fig S3B). *Gsdmc2-4* were robustly induced at the transcript and protein level following IL-4 treatment (Fig. 3D and 3E). As some of these early studies were underway in our lab, other groups also reported similar results, showing that *Gsdmc2-4* are downstream of type 2 immune signals and may play a role in protection against helminth infections^17,18^. Further treatment of organoids with cytokines only associated with Type 2 immune responses, including IL-4, IL-5, IL-9, IL-13, and IL-31 revealed that again, only IL-4 and IL-13 were able to induce *Gsdmc2-4* (Fig. S3C).

Given that type 2 immunity is regulated by the transcription factor STAT6, we next tested the necessity of STAT6 for Gasdermin C induction and confirmed that IL-4 and IL-13 induce STAT6 phosphorylation and led to an increase in GSDMC levels in intestinal organoids (Fig. 3F). Using WT or *Stat6^-/-^* mice^30^, we showed that ablation of STAT6 precluded Gasdermin C induction following IL-4 and IL-13 treatment in small intestinal organoids (Fig. 3G). Furthermore, basal expression of GSDMC2/3 protein in small intestinal crypts and villi was only partially diminished in *Stat6^-/-^*animals compared to WT mice (Fig 3H), which may indicate a more complex transcriptional regulation of Gasdermin C *in vivo*.

### The *Gsdmc* locus is responsive to microbial presence

As previously discussed, Gasdermin C exhibits spatially distinct patterns of expression. Both transcript and protein are enriched in the distal end of the gastrointestinal tract, with the highest expression in the colon. This led us to the idea that Gasdermin C may be regulated by microbial presence and load. To test this hypothesis, we obtained germ-free mice (GF) and compared levels of *Gsmdc2-4* transcripts and GSDMC2/3 protein levels between GF and conventional mice (CONV). We found that both transcript and protein levels were downregulated in the upper region of intestinal crypts and villi from GF mice (Fig. 4A-D). Interestingly, recolonization of germ-free mice (RC), using bedding from conventional mice, increased RNA and protein levels of Gasdermin C 2-4 in the small intestine (Fig. 4A and B). Surprisingly, *Gsdmc2-4* transcript and protein levels were nearly identical in colons of germ-free compared to conventional mice (Fig. S4A), suggesting a different mechanism of regulation.

**Figure 4:**
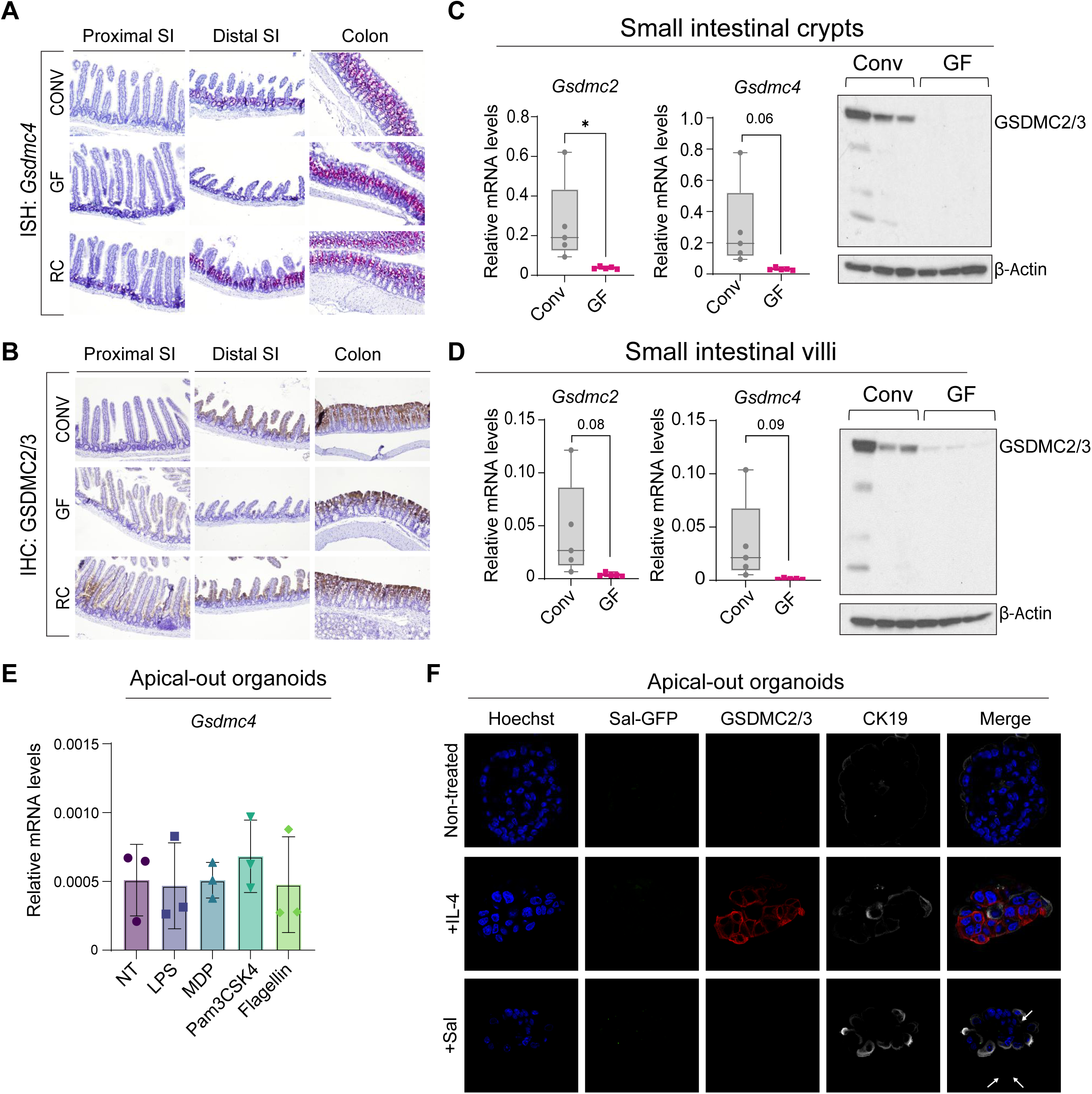
Microbial cues influence Gasdermin C2-4 distribution in GI. (A) RNAscope *in situ* hybridization staining of *Gsdmc4* in proximal and distal regions of mouse small intestine and in whole colon from conventional (CONV) mice, germ-free (GF) mice, and germ-free mice recolonized using bedding from conventional mice (RC) (n=3 mice per condition). (B) IHC staining of GSDMC2/3 in proximal and distal regions of mouse SI and in whole colon from conventional mice, germ-free mice, and germ-free mice recolonized using bedding from conventional mice (n=3 per condition). (C) SI crypts and (D) SI villi from conventional mice (CONV) and from germ-free mice were collected and used for RT-qPCR analysis for *Gsdmc2 and Gsdmc4* and western blot analysis for GSDMC2/3 (n=5-6 mice per condition). Representative samples are shown. (E) SI organoids were cultured for 3 days in WRN media, inverted to apical out organoids and grown in WRN media an additional 1-2 days. After inverting, organoids were treated with indicated TLR agonists for 24hrs before harvesting for RT-qPCR analysis of *Gsdmc4* (n=3 biological replicates). (F) Apical-out organoids were generated as described. After inverting, organoids were kept in suspension in WRN media for 24hrs, treated with IL-4 for 24hrs where indicated, then infected with GFP-Salmonella for 1hr. Fixed organoids were stained for GFP (green), GSDMC2/3 (red), and CK19 (white). Data are presented as mean ± SD. *p < 0.05.

To investigate whether *Gasdermin C* expression could be induced by specific bacterial signals, we generated apical-out organoids as described in *Co et al.*^31^ (Fig. S4B). These “inside-out” organoids have inverted polarity, allowing for easier access to the apical surface of epithelial cells, and providing a better representation of luminal interactions that may occur between epithelial cells and microorganisms. We treated murine small intestinal apical-out organoids with either bacterial components or other Toll-like receptor (TLR) agonists, such as LPS, MDP, PAM3CSK4, or flagellin for 24 hours. None of these compounds were able to induce *Gsdmc4* (Fig. 4E). To test if direct bacterial infection could induce Gasdermin C, we infected apical out organoids with GFP-tagged *Salmonella enterica* serovar Typhimurium (S. Typhimurium) prior to fixing and staining, or lysing cells. The infection was unable to induce expression of GSDMC2/3 protein (Fig. 4F and S4C).

Given that basal levels of *Gasdermin C* expression in our wild-type mice appeared higher than in mice from other recently published studies^17, 32^, this led us to seek other potential microbial factors that may be regulating *Gsdmc2-4* expression in our conventionally housed mice, including commensal microorganisms that may vary across housing facilities. C57BL/6J mice that were purchased from the Jackson Laboratory’s (JAX) maximum barrier level had lower expression of *Gsdmc2-4* transcript and protein in small intestinal crypts and villi than mice bred and raised in-house (OSU) (Fig. 5A-D). Again, we noted similar *Gsdmc2-4* expression patterns increasing from proximal to distal regions of the intestine in our mice, with JAX mice having substantially lower transcript (Fig. 5A) and protein levels overall (Fig. 5B-D). The colon did not exhibit much difference in *Gsdmc2-4* expression between mice from different facilities (Fig. S5A and S5B).

**Figure 5:**
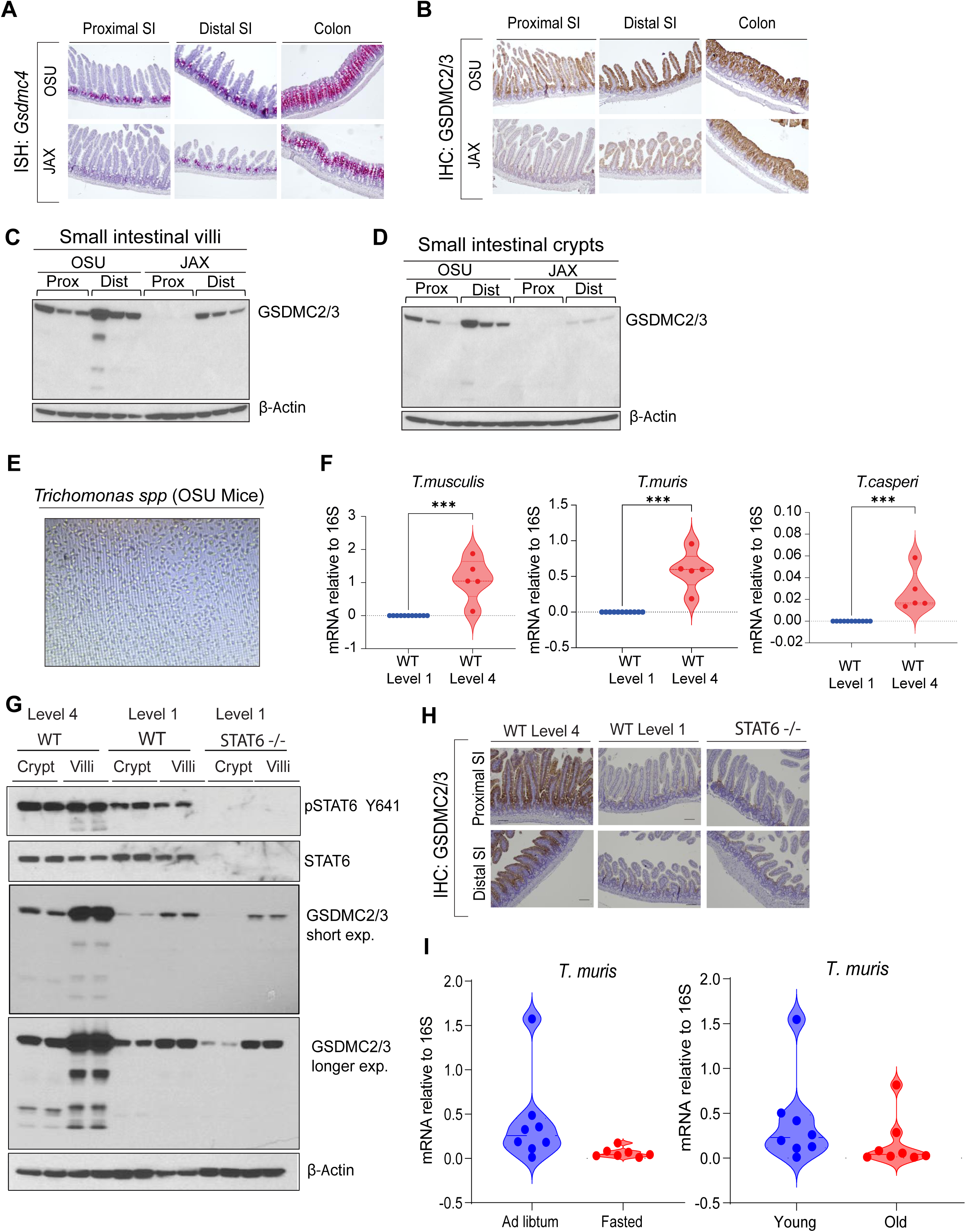
*Tritrichomonas* presence upregulates Gasdermin C2-4 expression through type 2 immunity. (A) RNAscope *in situ* hybridization staining of *Gsdmc4* and (B) IHC staining of GSDMC2/3 in proximal and distal regions of the mouse small intestine and in whole colon from mice bred and housed at OSU and mice purchased from the Jackson Laboratory’s maximum barrier level (JAX) (n=3 mice per condition). (C) Lysates from SI villi and (D) SI crypts from mice bred and housed at OSU (n=5-6 mice) and from mice purchased from the Jackson Laboratory’s maximum barrier level (n=3 mice) were collected and immunoblotted for GSDMC2/3 or (C-F) for RT-qPCR analysis. Representative samples are shown. (E) Bright-field scan of *Tritrichomonas spp.* found in cecal contents of OSU mice. (F) Fecal samples were collected from the cecum of WT Barrier Level 1 (n=11 mice) or Level 4 (n=5 mice), and microbial DNA was obtained for RT-qPCR analysis for *Tritrichomonas spp*. (G) Lysates from SI crypts and villi were collected from WT and *Stat6*^-/-^ (n=4 mice per condition) housed at either Barrier Level 1 or 4 for immunoblotting (H) IHC staining of GSDMC2/3 in proximal and distal regions of mouse small intestine collected from WT and *Stat6^- /-^* at either barrier level 1 or 4 (n=4 mice pre condition). Representative samples are shown. (I) Fecal samples were collected from the small intestines of *ad libitum* and 24hr fasted mice or from young (3-6 months) and old (22-30 months) mice and microbial DNA was obtained for RT-qPCR analysis of *Tritrichomonas muris* (n= 7-8 mice per condition). Data are presented as mean ± SD. ***p<0.001.

We next checked cecal contents and found that the cecal contents of OSU conventional mice were populated with the commensal protist *Tritrichomonas muris (T. muris)* (Fig. 5E), which was absent in the JAX mice. We then rederived our conventional mice to a Barrier Level 1 through cross-fostering which eliminates protist presence. High relative expression of *Tritrichomonas spp.* was confirmed in our standard mice (WT Level 4) compared to the re-derived mice (WT Level 1) (Fig. 5F). Additionally, under cleaner Level 1 housing conditions, GSDMC2/3 protein levels were found to be decreased compared to Level 4 conditions (Fig 5G and S5C). However, the absence of commensal protists did not completely abrogate Gasdermin C levels.

We next asked whether loss of STAT6 would affect *Gsdmc* levels *in vivo*. Although basal levels of GSDMC2/3 were lower in WT Level 1 mice compared to WT Level 4 mice, genetic ablation of *Stat6* under Level 1 conditions did not completely abolish GSDMC2/3 levels in villi, suggesting additional transcriptional regulation for baseline levels of *Gsdmc* expression (Fig. 5G, 5H and S5C). Interestingly, the observed reduction of *Tritrichomonas muris* during nutrient deprivation and in aged mice (Fig. 5I and S5D) suggests that loss of this protist may diminish Type 2 immune signaling, thereby contributing to the age- and nutrient-associated decline in *Gasdermin C* expression.

### Loss of *Gsdm1-4* exacerbates DSS-induced colitis in mice

We next generated *Gsmdc1-4* knockout mice *(Gsdmc^-/-^*, see Methods) and confirmed the absence of Gasdermin C transcripts and proteins (Fig. S6A and S6B). As no overt phenotypes were observed in the small intestine or colon at baseline, we next asked whether GSDMC contributes to epithelial protection during injury. To test this, we challenged WT and *Gsdmc^-/-^* mice maintained under level 4 (protist-colonized) housing, with 3.5% DSS for 7 days (Fig. 6A) and monitored body weight changes and signs of intestinal inflammation. Although the percent body weight loss (Fig. 6B), colonoscopy results (Fig. S6C), and small intestinal length (Fig.S6D) were comparable between DSS treated WT and *Gsdmc*^-/-^ mice, the colon length of the *Gsdmc*^-/-^ mice decreased significantly (Fig. 6C), which is the region most affected by DSS. Hematoxylin and Eosin staining and analysis showed that *Gsdmc* ^-/-^ mice had increased immune cell infiltration and crypt erosion compared to the WT mice (Fig. 6D and 6E). Given that loss of mucus-producing cells increases with DSS exposure, we stained tissues with Alcian blue, marking mucus producing cells, and observed that DSS treated *Gsdmc*^-/-^ mice had significantly fewer goblet cells compared WT, DSS treated counterparts (Fig. 6E, lower panel). Further analysis by a board-certified veterinary pathologist confirmed that *Gsdmc*^-/-^ mice had more extensive inflammation, mucosal loss with severe loss of crypts and mucosal epithelial erosion compared to WT counterparts (Figs. 6F and S6C). Altogether, these findings suggest that GSDMC may play an important role in preserving gut barrier integrity and maintaining mucus production in the intestinal epithelium following epithelial damage.

**Figure 6:**
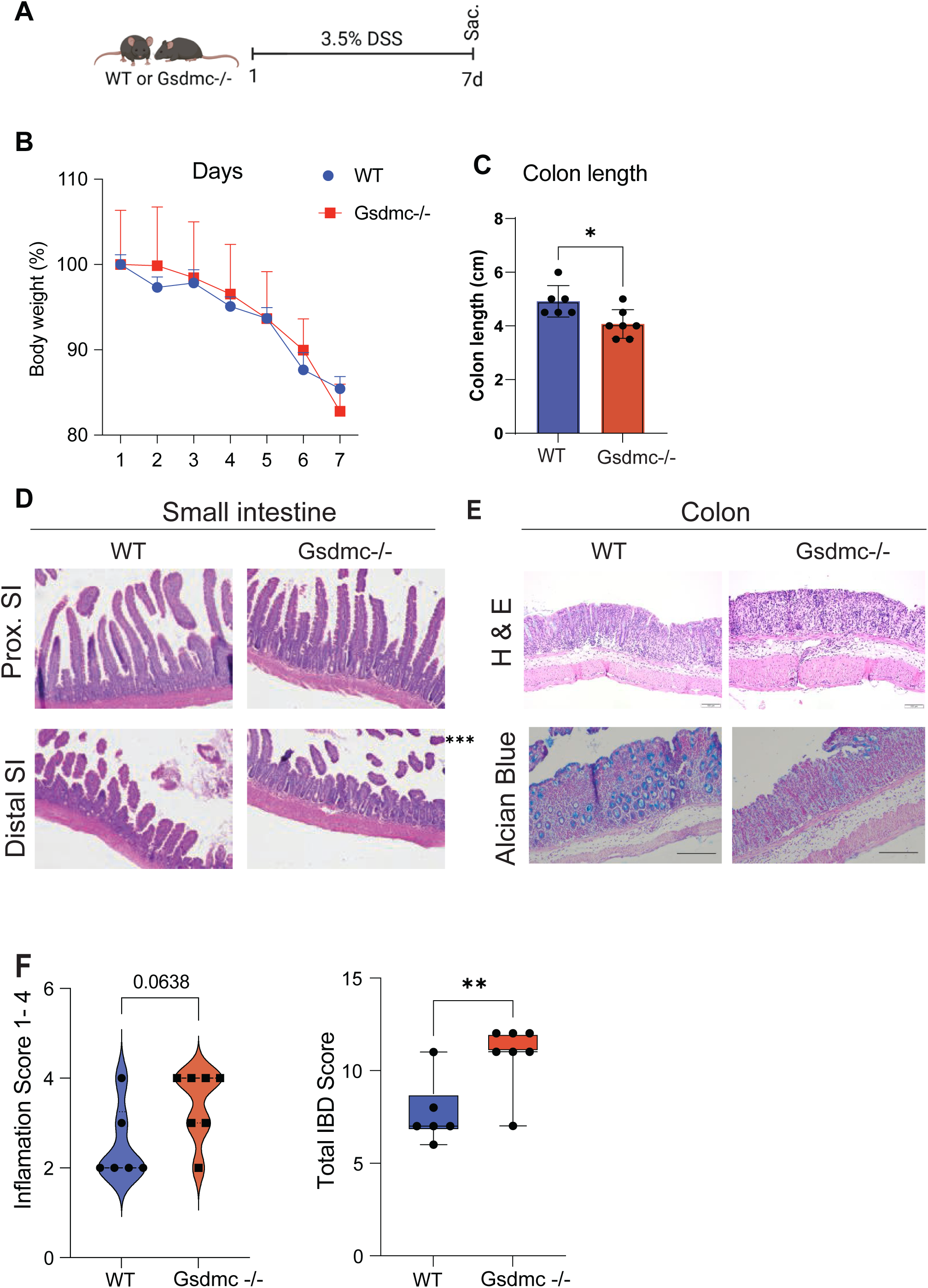
Loss of *Gsdm1-4* exacerbates DSS-induced colitis and tissue repair in mice. (A) Schematic of the DSS colitis experiment in protist-colonized WT (n=7) and *Gsdmc^-/-^* (n=8) mice. (B) Relative percent change in body weight between WT and *Gsdmc*^-/-^ mice. (C) Colon length of WT and *Gsdmc*^-/-^ mice subjected to DSS. (D) H&E staining of proximal and distal regions of the mouse small intestine of WT and *Gsdmc^-/-^* mice exposed to DSS. (E) Representative H&E-stained images (upper panel) and Alcian blue (lower panel) staining of colons from WT and *Gsdmc^-/-^* mice exposed to DSS. (F) Colitis severity scores as determined by histological analysis in WT and *Gsdmc^-/-^* mice exposed to DSS. Mice from both genotypes exhibit mucosal and submucosal inflammation, with inflammation and loss of crypts exacerbated in *Gsdmc^-/-^* mice. Data are presented as mean ± SD. **p<0.01.

## DISCUSSION

The intestine is a site of dynamic communication in which epithelial cells, immune cells, and the microorganisms residing in the intestinal lumen all engage in crosstalk to coordinate host homeostasis. The Gasdermin family of proteins, known for their ability to induce pyroptosis, are expressed across epithelial and mucosal surfaces in the body and is found in several types of immune cells^33^. Much of the work on these proteins has focused on signals that lead to their cleavage by caspases and subsequent induction of pyroptosis, yet little is known about conditions that affect their expression. Even less is known about GSDMC, which is thought to be selectively expressed in the gastrointestinal tract and epidermis^34,35^. Our study reveals a previously unappreciated layer of intestinal regulation, showing that GSDMC integrates signals from nutrients, microbes, and Type 2 immunity to dynamically shape epithelial function and gut repair across the lifespan.

We first observed that murine *Gasdermin C* expression (primarily *Gsdmc2-4*) increases from the proximal to the distal end of the gastrointestinal tract. We also observed that *Gsdmc2-4* transcripts are enriched in progenitor cells residing in the transit-amplifying zone in crypts of the small intestine, while the protein is produced in epithelial cells at the top of the crypt and across mature enterocytes up to the villi tips. Intriguingly, we rarely observed *Gsdmc* transcripts higher up the villus, contrasting with the protein localization primarily in villi, suggesting that the protein is likely produced in late enterocyte progenitors and stably maintained in more mature epithelial cells, as they move up the villus. This points to an interesting dynamic of transcription and translation carried out during specific stages of cell maturity and in precise zones of the crypt-villus axis. Others have previously reported that the upper crypt/lower villi zone is enriched in gene expression markers linked to barrier immunity and host defense function^36^, which Gasdermin C proteins are proposed to serve.

Closer evaluation of subcellular localization showed that Gasdermin C proteins co-localize with components of the terminal web^37^, a structure that can trap bacteria at the apical side of epithelial cells. This observation raises the possibility that GSDMC proteins may also participate in eliminating microbes attempting to breach the epithelial barrier. We and others have previously shown that aging is associated with impaired intestinal homeostasis and leads to diminished nutrient absorption, decreased immune function, and imbalance of the gut microbiome, compromising the gut barrier and epithelial regeneration^38^. Interestingly, we observed lower Gasdermin C abundance in aged mice (22-30mo), both at the transcript and protein level, whereas Gasdermin D levels remained unchanged between young and aged mice. These findings suggest that age-associated GSDMC decline may contribute to impaired epithelial regeneration, potentially linking altered epithelial-microbial-immune crosstalk to the deterioration of gut barrier integrity in aging.

Although the Gasdermin C proteins appeared to be remarkably stable, transcripts were highly responsive to changes in the intestinal environment. Nutrient deprivation, in the form of a 24-hour fast, nearly abolished *Gsdmc2-4* expression throughout the small intestine. Mice that underwent a 24-hour re-feed, following the fasting period, showed re-expression of *Gsdmc2-4* in the presence of nutrients. We predict that these changes stem due to lack of nutrients available to the protist and suppression of type 2 immune responses in the epithelium. Given that GSDMC levels increase from proximal to distal in the small intestine, we hypothesized that bacterial load, or specific strains of bacteria, may be necessary to induce *Gsdmc2-4*. However, exposure to several common bacterial components or TLR agonists, typically present in commensals added into apical-out organoid cultures did not induce GSDMC. Other studies have demonstrated that *Gsdmc* transcription is regulated downstream of IL-4/13-STAT6 signaling in the context of helminth infection^18^ and have observed low expression of *Gsdmc* transcripts in organoid culture^39^ prior to addition of type 2 cytokines. In parallel, we also found similar results with type 2 specific induction of *Gsdmc2-4* transcripts and GSDMC proteins in cultured organoids and further extended that work to show that STAT6 is needed for IL-4/13-mediated *Gsdmc* induction in intestinal organoids. However, STAT6 may not be the only transcription factor that upregulates Gsdmc *in vivo,* as basal levels of *Gsdmc* expression could still be detected in our *Stat6* KO mice. It is worth noting that while *Gsdmc* expression was not directly induced by microbial cues or non-type 2 cytokines, these signals may still play a role in directing Gsdmc protein cleavage and/or pore formation in intestinal epithelial cells^40^.

Separately, examination of the cecal contents of our mice revealed that they are colonized with non-pathogenic commensal protist species, such as *Tritrichomonas muris*. These protists are known to activate type 2 immune circuits in the gut^22^. This fits well with our initial findings that type 2 immune cytokines can robustly induce *Gsdmc* expression in small intestinal organoid cultures that are sterile and that Godecs were downregulated in NOD-SCID mice. Furthermore, we found that the levels of *T. muris* can change in response to nutrient availability and aging, which may partially explain our initial observations that Gasdermin C levels can fluctuate in response to these two conditions. Collectively, these findings highlight the important role of nonpathogenic microbial communities in shaping and supporting intestinal immune responses. Based on our observations, we were able to see potential cleavage products in mouse samples in the absence of pathogenic infection. If they do indeed generate pores, this type of activity may be homeostatic and have a specific immune function feeding back to a heightened type 2 immune response in the commensal colonized mice. Moreover, our immunofluorescence or immunobiological analysis cannot distinguish between the full length and cleavage products, warranting future investigation into the role of these products. Additionally, future studies will be needed to determine how changes in microbial populations, particularly those occurring with aging or nutrient fluctuations, modulate crosstalk among immune regulatory pathways and impact gut immune homeostasis.

There have been several exciting and recent findings on the role of GSDMCs in intestinal homeostasis and repair from injury, both in the context of sterile injury and with infection. While some reports indicate that GSDMC may be dispensable for tissue recovery following DSS induced injury^39^, others have reported detrimental effects for epithelial cell specific GSDMC expression in driving DSS-mediated pathology^40^. In our study, global loss of *Gsdmc1-4* worsened DSS-induced pathology in protist-colonized mice, highlighting context dependent protective roles of GSDMCs in epithelial injury and integrity. Interestingly, a recent study also reported that m^6^A depletion can trigger *Gsdmc* transcript degradation, leading to mitochondrial dysfunction, while human GSDMC knockdown led to apoptosis in human colonic organoids^41^.

Given that we observed lower levels of Gasdermin C proteins in aged mice, this raises the possibility that age-related GSDMC decline may contribute to impaired gut regeneration and susceptibility to injury. Type 2 immunity influences various aspects of intestinal function, and thus it remains to be seen how heightened and prolonged type 2 immune responses, and their interactions with GSDMC, impact gut homeostasis, inflammatory bowel predisposition and colorectal cancer^42^. Collectively, these findings underscore context-dependent and multifaceted roles for GSDMC in intestinal homeostasis and repair and highlight the need for future studies examining its age-dependent, immune-modulated, and microbiome-influenced functions.

## RESOURCE AVAILABILITY

### STAR METHODS

- **KEY RESOUCES TABLE**
- **EXPERIMENTAL MODEL AND PARTICIPANT DETAILS**

#### Lead contact

Further information and requests for resources and reagents may be directed to and will be fulfilled by the lead contact, Maria Mihaylova;

#### Materials availability

Mouse and microbial strains used in this study will be made available upon request, addressed to the lead contact.

## ACKNOWLEDGMENTS

We thank Drs. Min Chen and Foued Amari for the generation of the mouse line at the Genetically Engineered Mouse Modeling Core of the College of Medicine and James Comprehensive Cancer Center, The Ohio State University. We thank members of the Mihaylova laboratory for further additional comments and editing. The work, and M.M.M, were supported in part by NIH R00AG054760, the Glenn Foundation and American Federation for Aging Research, Pew Biomedical Scholar Award, and start-up funds from the Ohio State Comprehensive Cancer Center. A.K. was supported in part by Pelotonia Graduate Fellowship. A.K and O.W were supported in part by NIH T32 GM141955. J.S was supported in part by Pelotonia Postdoctoral Fellowship and Cancer Prevention NIH T32 CA2291140-02. We thank the Comparative Pathology & Digital Imaging Shared Resource at The Ohio State University Comprehensive Cancer Center, Columbus, OH for histology support and histopathological evaluation. Images presented in this manuscript were generated using the instruments and services at the Campus Microscopy and Imaging Facility, The Ohio State University (RRID:SCR_025078). These resources are supported in part by grant P30 CA016058, National Cancer Institute, Bethesda, MD.

## AUTHOR CONTRIBUTIONS

A.K. and M.M.M. conceptualized the study with the help of A.A. A.K., A.A., carried out majority of experiments with the assistance of K.B.A.C., M.F., J.S., and K.G. M. F. maintained the mouse colony and aided in mouse related studies. V.C. led the generation of Gasdermin C knockout mice. A.S. and B.A. guided and assisted with Salmonella experiments. R.K., P.R. and A.F. provided reagents and consulted on immune related studies. S.E.K. contributed to DSS colitis investigation and histopathologic scoring. A.K, and M.M.M. wrote the manuscript with the help of A.A. M.M.M. supervised the project. All authors reviewed the final manuscript.

## STAR METHODS

### EXPERIMENTAL MODEL

#### Animal studies

Germ-free C57BL/6NCrl mice were purchased from Charles River Laboratories (strain no. 574) and euthanized within 24 hours of arrival. C57BL/6J mice were purchased from the Jackson Laboratory (strain no. 000664) from a maximum barrier level. C57BL/6J mice were also bred in-house and periodically refreshed with new C57BL/6 mice purchased from the Jackson Laboratory. NOD-SCID mice (strain no. 001303), *Stat6^-/-^* mice (strain no. 005977), and Cmv-Cre mice (006054) were purchased from the Jackson Laboratory.

For aging experiments, young mice were between 3 to 6 months of age and old mice were between 20 and 28 months at the time of the experiments. For fasting experiments, singly housed mice were moved to a new cage and food was completely removed beginning at 9AM. Mice were euthanized 24 hours later or given food for an additional 24 hours before sacrifice. *Ad libitum* mice were also moved to a new cage and singly housed but given free access to chow for the duration of the experiment. Mice were age- and sex-matched. All animals were housed at The Ohio State University and studies were performed in accordance with the Institutional Animal Care and Use Committee guidelines at OSU.

#### Generation of *Gsdmc* KO animals

To generate *Gsdmc* null mice, CRISPR/Cas9 technology was used to target and insert *loxP* sites flanking the entire *Gsdmc* gene to make a conditional allele (*fl*); recombination between the two *loxP* sites will completely remove *Gsdmc1-4*. Two founder females had at least flanking *loxP* sites *cis* on one chromosome, and one bred to produce some offspring that were *Gsdmc^fl/+^*. This work was performed by the Genetically Engineered Mouse Modeling Core (GEMMC) at OSU. *Gsdmc^fl/+^* mouse was bred to *CMV-Cre^+^* mice^43^ to induce germ-line recombination between the *loxP* sites to create the null allele. *Gsdmc^+/-^* mice were backcrossed at least four generations to C57BL/6J to backcross and separate the *Cre* allele, then intercrossed to generate *Gsdmc^-/-^* mice. Genotyping was performed on ear biopsies with the following primer sets: 5’ loxP site used *Gsdmc* F and *Gsdmc* 5’ R to produce 514bp WT and 548bp mutant products; 3’ loxP site used *Gsdmc* 3’ F and *Gsdmc* R to produce 349bp WT and 383bp mutant products; and recombination and removal of the *Gsdmc* gene used *Gsdmc* F and *Gsdmc* R to produce only a 484bp deletion product. All animals were housed at the Ohio State University and in accordance with Institutional Animal Care and Use Committee guidelines at OSU. Primers for genotyping the *Gsdmc* loci are listed in the Key Resource Table.

### METHOD DETAILS

#### Crypt isolation and organoid cultures

Small intestinal crypts were isolated as described previously^44^. Isolated crypts were resuspended and adjusted to an approximate density of 5-10 crypts per µl in small intestinal crypt culture media, which consists of Advanced DMEM F12 (Gibco), 2mM L-Glutamine (Gibco), 100U/ml Penicillin (Gibco), 1000ug/ml streptomycin (Gibco), 200ng/ml Noggin (Peprotech), 50ng/ml R-spondin, 1X B27 (Life Technologies), 3µM CHIR99021 (Stem Cell Technologies), 1X N2 (Life Technologies), 10µM Y-27632 dihydrochloride monohydrate (Sigma Aldrich), 1µM *N*- acetyl-L-cysteine (Sigma Aldrich), and 15ng/ml EGF (Peprotech). Isolated crypts were plated in 25µl droplets containing 70% Matrigel (Corning) and 30% crypts in crypt culture media in 48-well plates (Corning) and allowed to solidify in a 37°C incubator for 15min before 250µl of crypt culture media was overlaid on the Matrigel droplets. The organoid cultures were maintained in 37°C humidified incubators containing 5% CO_2_, and media was refreshed every 2-3 days.

Where applicable, villi were also collected after filtering out crypts. Crypts and villi to be used for RT-qPCR or western blot analysis were either collected from the entire length of the small intestine, or the small intestine was cut in half and designated as proximal and distal before proceeding with the isolation procedure. To isolate colon crypts, the colon was removed, flushed with PBS, and opened longitudinally. Mucus was physically removed from the tissue using a microscope slide, and the tissue was washed in PBS in a 50mL conical tube by inversion, replacing the PBS 3x. The tissue was then minced into approximately 5mm size pieces and transferred to a 15ml conical tube with PBS plus EDTA (5mM). Tissue pieces were pipetted up and down to disrupt crypts, allowed to settle, and the solution was replaced with fresh PBS plus EDTA. The tissues were then incubated at 4°C for 15min.

Colon crypt base media was prepared, adapted from O’Rourke et al. 2016^45^, consisting of Advanced DMEM F12 (Gibco), 2mM L-Glutamine (Gibco), 100U/ml Penicillin (Gibco), 1000µg/ml streptomycin (Gibco), 1mM *N*-acetyl-L-cysteine (Sigma Aldrich), 10mM HEPES (Sigma Aldrich), and 10mM nicotinamide (Sigma Aldrich). The PBS/EDTA solution was removed, tissues were washed with cold PBS twice, and the PBS was removed as far as possible before 3mls of pre-warmed digestion media was added to each tube, containing colon crypt base media supplemented with 500U of Collagenase IV (Worthington Biochem). The tissues were incubated in a 37°C bead bath for 30min. To quench the reaction, 10mls of cold PBS was added to each tube and samples were inverted, then pipetted up and down approximately 10x to dislodge crypts and the solution was transferred to a new 15ml conical tube. The process was repeated two more times. Crypt fractions were combined and filtered through a 70µm mesh into a 50ml conical tube for downstream analyses.

#### Cytokine treatments, inhibitor treatments

Small intestinal organoids were generated as described above and various cytokines were added into the media on day 4 for 48 hours. Organoids were harvested on day 6. Cytokines used were murine IL-4 (5ng/ml or 10ng/ml), IL-5 (20ng/ml), IL-6 (40ng/ml), IL-9 (20ng/ml), IL-10 (10ng/ml), IL-13 (10ng/ml or 20ng/ml), IL-17 (20ng/ml), IL-22 (5ng/ml), IL-31 (20ng/ml), IFNγ (20ng/ml), and IFNλ (500ng/ml) (all recombinant cytokines were purchased from Peprotech). Organoids were then harvested for RNA extraction.

#### Apical out organoids

Small intestinal organoids were generated as described above and cultured in WRN-conditioned media for 3 days. Organoids were dislodged by resuspending and scraping both organoids in Matrigel in 5mM EDTA in PBS, then incubating for 1.5hrs on a rotator at 4°C. The EDTA-PBS solution was replaced after 1 hour. The organoids were allowed to settle by gravity, the supernatant removed, and the organoids were resuspended in 750µl WRN conditioned media per well of a 24 well Ultra Low Attachment plate. Media was changed 24hrs after the inversion process to remove dead cells by collecting organoids and allowing them to settle by gravity, removing the supernatant, and resuspending in fresh media. For treatments, bacterial compounds were added into the media for 24 hours. Compounds included: LPS (1μg/ml), MDP (10μg/ml), PAM3CSK4 (500ng/ml), and flagellin (500ng/ml). For bacterial infection, organoids were cultured for 48 hours after the inversion process with one media change. Twenty-four hours prior to infection, IL-4 was added to the media (5ng/ml). Organoids were infected with GFP-tagged *Salmonella typhimurium* for 1 hour in a 37°C incubator with 5% CO_2_, then collected and washed with PBS, and either fixed in 4% paraformaldehyde for whole mount staining or lysed in RIPA buffer.

#### Protist Quantification

Fecal samples were collected from the small intestine or cecum, and microbial DNA was extracted using ZymoBIOMICS DNA/RNA Miniprep Kit following the manufacturer’s protocol. The microbial DNA was used for RT-qPCR analysis to assess the level of *Tritrichomonas spp* relative to *16S* (primers are listed in Table 1).

#### RNA Isolation and RT-qPCR

Total RNA was extracted from tissue or organoids using TRI Reagent (Life Technologies) and purified using the RNeasy Mini Kit (Qiagen) following the manufacturer’s instructions. RNA was reverse transcribed using cDNA Supermix qScript (QuantaBio) and then diluted 1:3 prior to amplification with SYBR Green Supermix (Bio-Rad). Primers are listed in Table 1.

#### *In situ* hybridization

Small intestine and colon tissues were fixed in 10% formalin, paraffin-embedded, and sectioned. Single-molecule *in situ* hybridization was performed using the RNAscope 2.5 HD Detection Kit-Red (Advanced Cell Diagnostics) according to the manufacturer’s protocol. Sections were incubated with RNAscope Target Retrieval Reagents for 15min and RNAscope Protease Plus Reagent for 30min. Probes used in this study are as follows: *Mm-Gsdmc4* and *Mm-Gsdmd*.

#### Histology, Immunohistochemistry, and Immunofluorescence

Small intestine and colon tissues were fixed in 10% formalin and paraffin embedded. Sectioned tissues were subjected to antigen retrieval using Borg Decloaker RTU Solution (Biocare Medical) and incubated with 0.3% H_2_O_2_. Slides then underwent blocking with 5% normal donkey serum (Jackson ImmunoResearch) in TBST for 30min, followed by subsequent 15min incubations in 5% donkey serum plus avidin/biotin blocking kit (Vector) solutions. <u>Antibodies used</u> <u>for immunohistochemistry:</u> rabbit anti-Gsdmc2/3 (1:2000, Abcam ab229896). Biotin-conjugated donkey anti-rabbit secondary antibody (1:500, Jackson ImmunoResearch) was used with the Vectastain Elite ABC Immunoperoxidase detection kit (Vector Labs) and Dako Liquid DAB+ Substrate (Dako) for visualization. Antibody dilutions for immunohistochemistry were made in Common Antibody Diluent (BioGenex). Slides were counterstained with 50% hematoxylin and mounted with Cytoseal XYL (Thermo). Slides for immunofluorescence underwent identical antigen retrieval and peroxide incubations prior to blocking and permeabilization using 5% normal donkey serum in 0.2% Triton X-100 in PBS. <u>Antibodies used for immunofluorescence:</u> rabbit anti- GSDMC2/3 (1:400, Abcam ab229896), rat anti-keratin 19 (1:9) Developmental Studies Hybridoma Bank, University of Iowa, TROMA-III), mouse anti-E-cadherin (1:200, BD Biosciences BDB610182), mouse anti-villin (1:200, Santa Cruz sc58897). Alexa Fluor Donkey anti-rabbit 488, Alexa Fluor Donkey anti-mouse 546, Alexa Fluor Donkey anti-rat 647 secondary antibodies were used throughout this study (1:400, Thermo) and Hoechst (1ug/ml in PBS, Thermo).

#### Organoid embedding and staining

For whole mount staining, organoids were fixed with 4% paraformaldehyde for 20min at room temperature and permeabilized and blocked with 2% normal donkey serum in 0.2% Triton X-100. Organoids were incubated with primary antibodies diluted in PBST and 2% donkey serum. <u>Antibodies used:</u> rabbit anti-GSDMC2/3 (1:400, Abcam ab229896). Organoids were washed and then incubated with secondary antibody Alexa Fluor Donkey anti-rabbit 488 (1:400) for 1 hour at room temperature, followed by incubation with Hoechst solution (1ug/ml in PBS). For embedding and sectioning, organoids were again fixed with 4% paraformaldehyde for 20min at room temperature, then stained with methylene blue (2% in PBS) and incubated in 30% sucrose overnight at 4°C prior to embedding in optimal cutting temperature compound. Sectioned organoids were washed with PBS and used for immunofluorescence protocol. <u>Antibodies used:</u> rabbit anti-GSDMC2/3 (1:400, Abcam ab229896) and mouse anti-E-cadherin (1:200, BD Biosciences BDB610182). Secondary antibodies used were Alexa Fluor Donkey anti-rabbit 488 and Alexa Fluor Donkey anti-mouse 546.

#### Immunoblotting

Isolated crypts and villi were lysed in RIPA buffer (50mM HEPES pH 7.4, 40mM NaCl, 2mM EDTA, 1.5mM sodium orthovanadate, 50mM NaF, 10mM sodium pyrophosphate, 10mM sodium β-glycerophosphate, 0.1% SDS, 1% sodium deoxycholate, 1% Triton) supplemented with cOmplete Protease Inhibitor Cocktail (Roche) and PhosSTOP phosphatase inhibitors (Roche). Lysates were normalized and equal amounts loaded onto Novex precast gels (Life Technologies). <u>Antibodies used:</u> rabbit anti-GSDMC2/3 (1:1000, Abcam ab229896), mouse anti-βeta-Actin (1:2000, Santa Cruz sc-47778), rabbit anti-STAT6 (1:1000, Abcam ab32520), rabbit anti-pSTAT6 Y641 (1:500, Abcam ab263947).

#### DSS Colitis Model

WT and *Gsdmc^-/-^* mice were supplemented with 3.5% DSS (36–50kDa) in their drinking water for 7 days. Mice were weighed and monitored daily for dehydration, weight loss, and blood in the stools. Mice were sacrificed on day 7, and small intestine and colon tissues were collected and fixed in formalin for histological analysis, then analyzed by board certified veterinary pathologist (SEK).

#### DSS Histopathology and Statistics

The gastrointestinal tract was isolated, and the colon was separated from the cecum. Samples of colon, cecum and occasionally mesenteric lymph nodes were placed into 10% neutral buffered formalin. The colon was prepared in a ‘‘Swiss roll’’ technique, routine-processed for paraffin embedding and stained with Hematoxylin & Eosin (H&E). H&E-stained colon sections were evaluated by a board-certified veterinary pathologist (SEK) blinded to the experimental groups. Colitis scoring was adapted from previously reported methods ^46,47^ and included scoring the entire colon for the severity of mucosal loss, mucosal epithelial hyperplasia, degree of inflammation and extent of pathology on a 0–4 scale. Scores were summed to calculate the total IBD score. An overall score was submitted for each section evaluated. To compare one variable between two groups, unpaired t-tests were used, or Mann-Whitney tests was used if the data were not normally distributed.

#### Data Analysis

Unless otherwise stated, most in vivo experiments were conducted with 4-8 mice and/or repeated three times. In the main text and figures, “n” represents biological replicates. For organoid- related experiments, at least 2-4 wells were pooled for Western blot analysis, with a minimum of 3-4 biological replicates. Statistical significance was assessed by primarily by Student’s t-test, unpaired unless otherwise indicated. No samples or animals were excluded from the analysis. Sex-matched young mice (3-6 months old) or aged mice (20-28 months old) were randomly assigned for *in vivo* studies. All animal experiments were approved by The Ohio State University Institutional Animal Care and Use Committee (IACUC).

---

**Supplemental Figure 1. Gasdermin C2-4 levels are regulated by nutrient availability in mice**

(A-D) Small intestinal crypts and villi, as well as colon crypts were collected from *ad libitum*, 24hr fasted, and 24hr refed mice. Gasdermin C protein levels were determined by western blot. *Gsdmc* expression was determined by RT-qPCR in colon crypts (E) and *Gsdmd* expression was determined by RT-qPCR in small intestinal crypts (F).

**Supplemental Figure 2. Aging suppresses Gasdermin C2-4 levels in mice**

(A-C) Proximal and distal small intestine crypts and whole colon crypts from young (3-6mo, n=5- 6) and old (22-30mo, n=5-6) mice were collected and expression of *Gsdmc2*, *Gsdmc3*, *Gsdmc4*, and *Gsdmd* genes were determined by RT-qPCR.

**Supplemental Figure 3. Gasdermin C2-4 expression is regulated by Type 2 immune signals**

(A) Colon crypts from WT (n=5-6) and NOD-SCID mice (n=4) were collected and used for RT-qPCR analysis for *Gsdmc2* and *Gsdmc4* and immunoblotted for GSDMC2/3. Representative samples are shown. (B) SI organoids were cultured for 4 days and treated with cytokines for 48hrs. Expression of *Cxcl9 and Isg15* genes were determined by RT-qPCR (n=5 biological replicates). (C) SI organoids were cultured for 4 days and treated with cytokines for 48hrs. Expression of *Gsdmc2*, *Gsdmc3*, and *Gsdmc4* genes were determined by RT-qPCR (n=5 biological replicates). Data are mean presented as ± SD. *p < 0.05, **p<0.01, and ***p<0.001.

**Supplemental Figure 4. Salmonella infection does not trigger Gasdermin C2-4 expression**

(A) Colon crypts from conventional mice (Conv) and germ-free (GF) mice were collected and used for RT-qPCR analysis (n= 5 per condition) for *Gsdmc2* and *Gsdmc4* and immunoblotted for GSDMC2/3. (B) Small intestine organoids were inverted to apical-out organoids. Fixed organoids were stained for Phalloidin (red). (C) After inverted organoids were established, they were treated with IL-4 for 24hrs as indicated and then infected with GFP-Salmonella for 1hr. Lysates were collected and immunoblotted for GSDMC2/3 and caspase-3 (n=3-6 biological replicates). Representative samples are shown.

**Supplemental Figure 5. Expression of *Gasdermin C2-4* in the colon is not dependent on *Tritrichomonas* status or STAT6**

(A) Lysates from colon crypts of mice bred and housed at OSU (n=5-6) and from mice purchased from the Jackson Laboratory’s (JAX) maximum barrier level (n=3) were collected and immunoblotted for GSDMC2/3 or (B) used for RT-qPCR analysis. Representative samples are shown. (C) IHC staining of GSDMC2/3 in mouse whole colon collected from WT (n=4) and *Stat6-/-* (n=4) mice at either Barrier Level 1 or 4. Representative samples are shown. (D) Fecal samples were collected from the small intestines of *ad libitum* (n=6), 24hr fasted (n=6), and 24hr refed (n=6) mice, and microbial DNA was obtained for RT-qPCR analysis for *Tritrichomonas muris*. Data are presented as mean ± SD. *p < 0.05, **p<0.01.

Supplemental Figure 6. Loss of *Gsdmc1-4* worsens intestinal inflammation in DSS-induced colitis mouse model

(A) Representative ISH images of *Gsdmc4* in non-treated, ad libitum WT and *Gsdmc^-/-^* mice. (B) Representative IHC images of GSDMC2/3 in non-treated, ad libitum WT and *Gsdmc^-/-^* mice. (C) Colonoscopy images and (D) small intestinal length of DSS treated WT and *Gsdmc^-/-^* mice. Pathology analysis of mucosal loss in colons of DSS treated WT and *Gsdmc^-/-^* mice (n=7-8 mice per condition). Data are presented as mean ± SD. ***p<0.001

