## Supplementary figures and images for "Gasdermin C links nutrient and immune signaling to protist-induced type 2 immunity and intestinal repair"

### Supplemental Figures

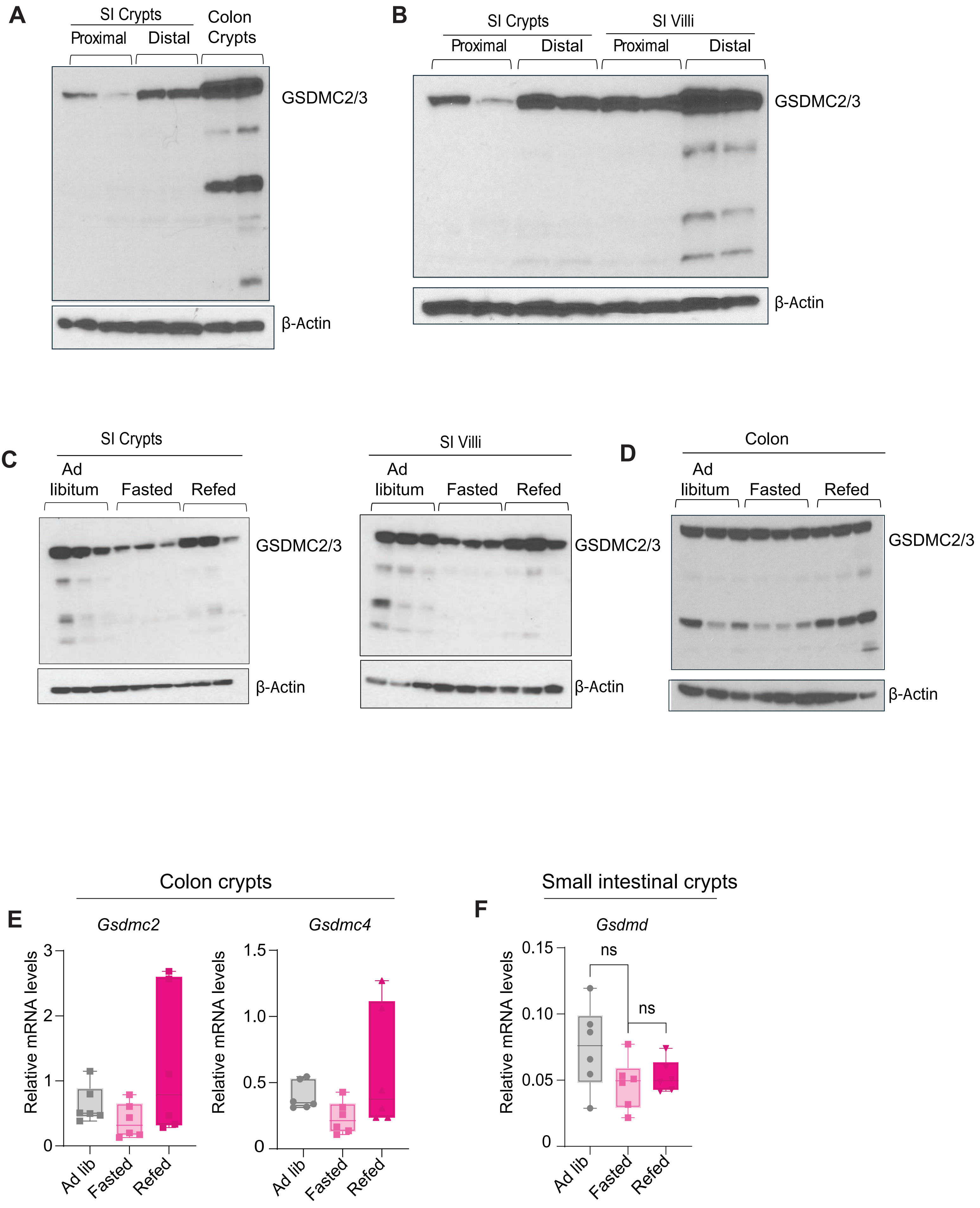

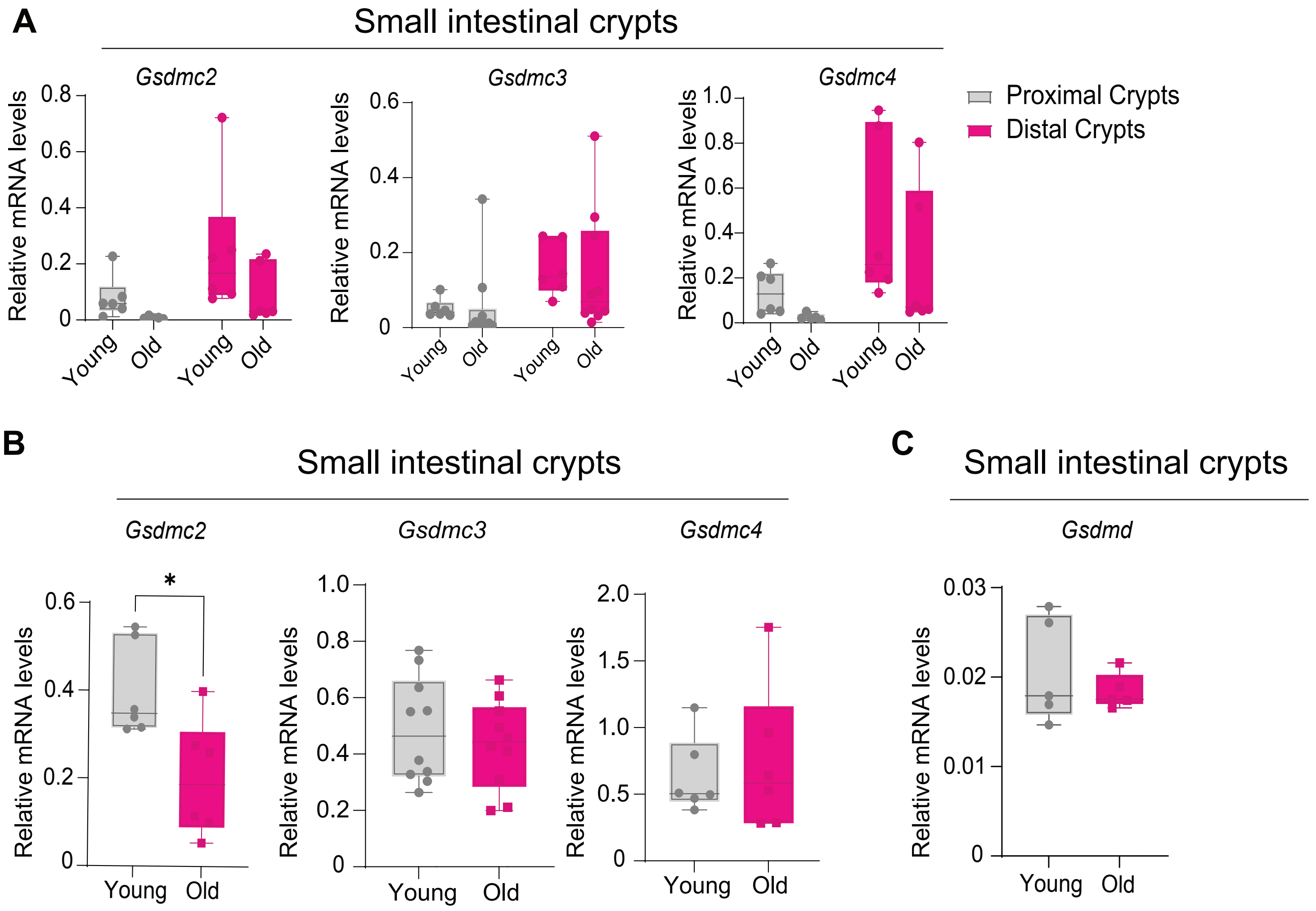

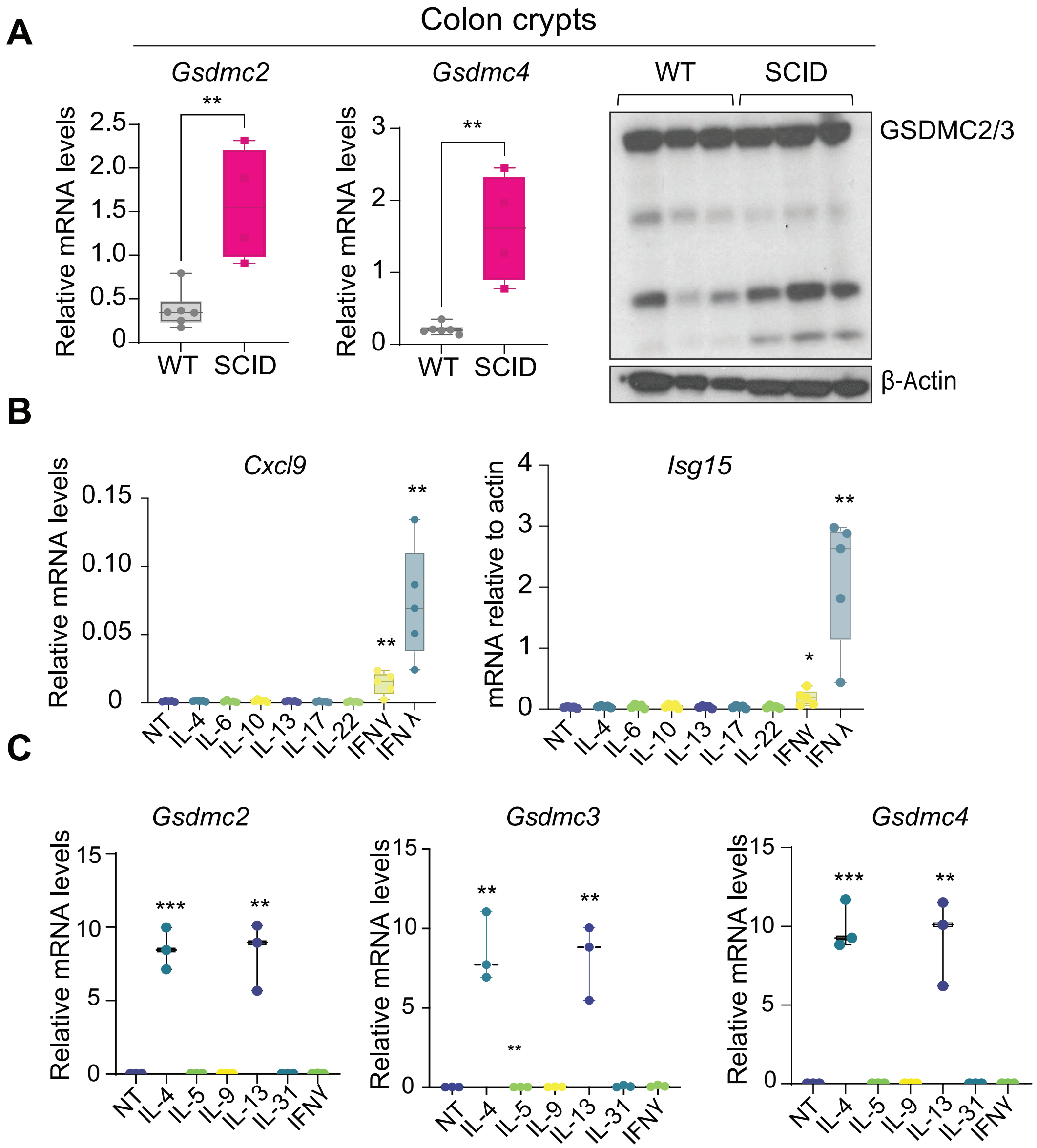

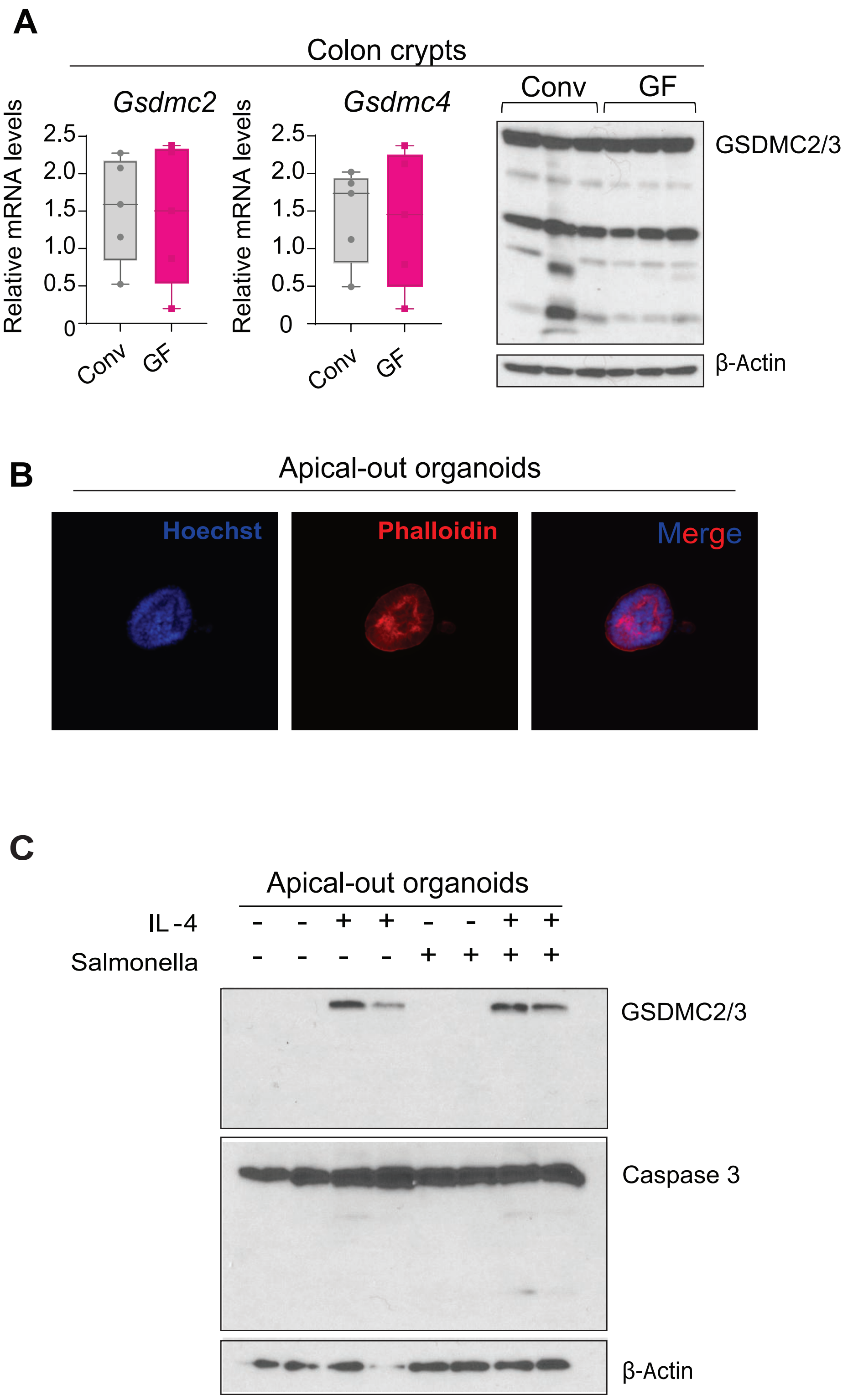

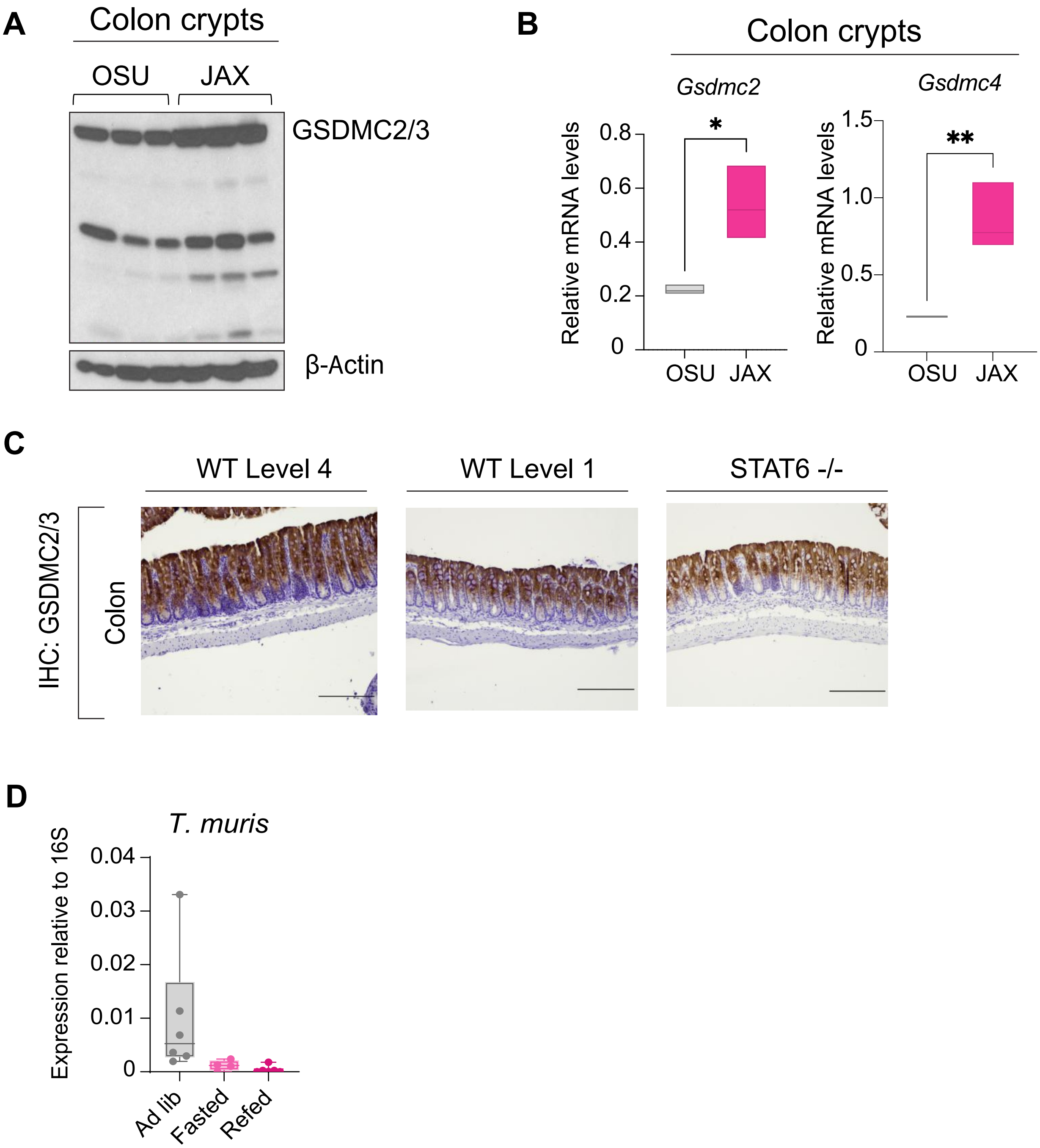

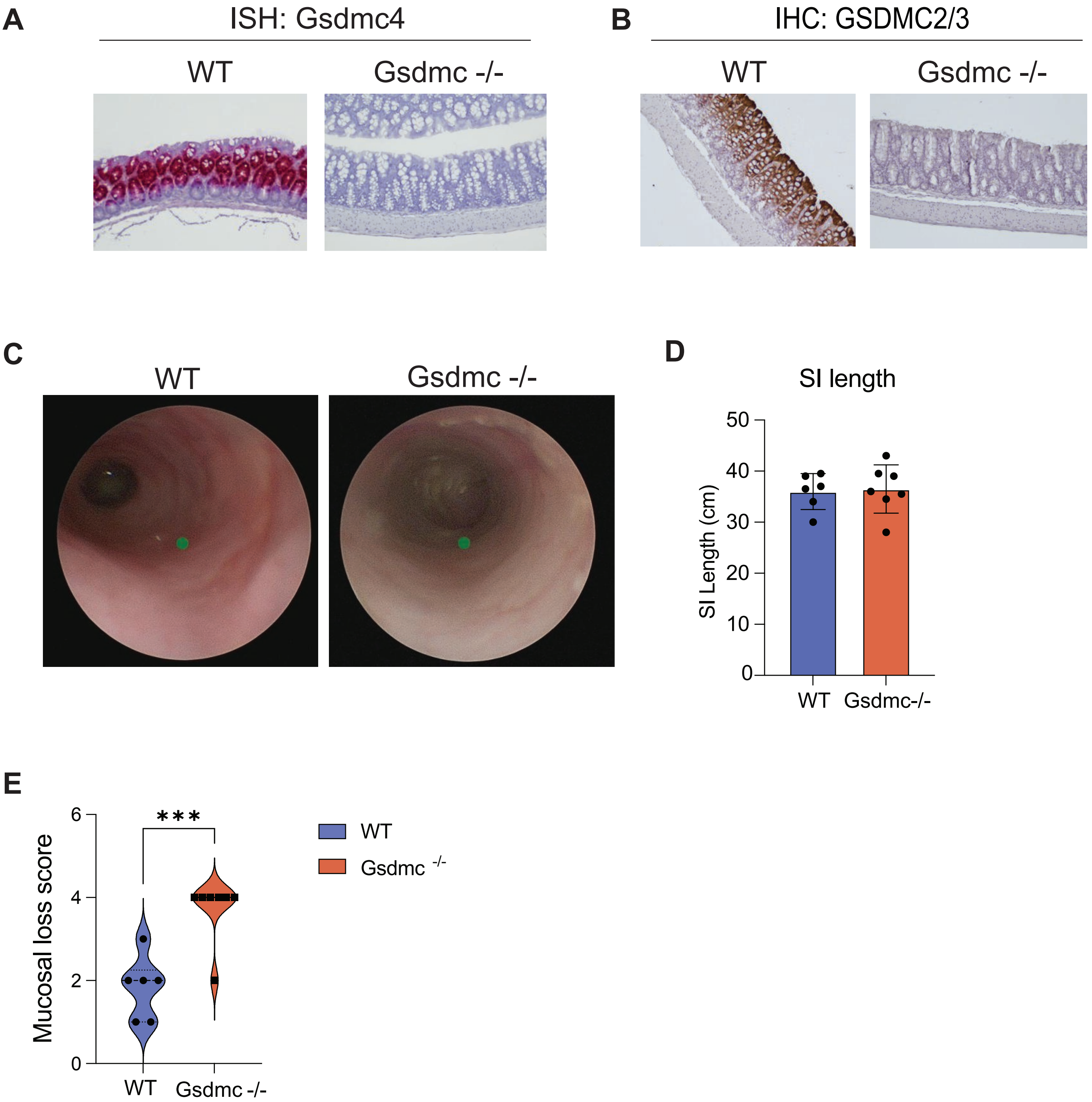
